# DBSCAN-SWA: an integrated tool for rapid prophage detection and annotation

**DOI:** 10.1101/2020.07.12.199018

**Authors:** Rui Gan, Fengxia Zhou, Yu Si, Han Yang, Chuangeng Chen, Jiqiu Wu, Fan Zhang, Zhiwei Huang

## Abstract

**Summary:** As an intracellular form of a bacteriophage in the bacterial host genome, a prophage is usually integrated into bacterial DNA with high specificity and contributes to horizontal gene transfer (HGT). Phage therapy has been widely applied, for example, using phages to kill bacteria to treat pathogenic and resistant bacterial infections. Therefore, it is necessary to develop effective tools for the fast and accurate identification of prophages. Here, we introduce DBSCAN-SWA, a command line software tool developed to predict prophage regions of bacterial genomes. DBSCAN-SWA runs faster than any previous tool. Importantly, it has great detection power based on analysis using 184 manually curated prophages, with a recall of 85% compared with Phage_Finder (63%), VirSorter (74%) and PHASTER (82%) for raw DNA sequences. DBSCAN-SWA also provides user-friendly visualizations including a circular prophage viewer and interactive DataTables.

**Availability and implementation:** DBSCAN-SWA is implemented in Python3 and is freely available under an open source GPLv2 license from https://github.com/HIT-ImmunologyLab/DBSCAN-SWA/.

## 1 Introduction

Bacteriophages are viruses that specifically infect their bacterial hosts. Passive replication of the phage genome relies on integration into the host’s chromosome and becoming a prophage (Panis et al., 2010). Prophages coexist and evolve with bacteria, influencing the entire ecological environment. Recently, phage therapy, defined as using phages to treat bacterial infections, has also been greatly emphasized. Therefore, the identification of prophages in their host genomes become critical not only for understanding their biological mechanisms but also for developing therapeutic strategies.

Several tools have been developed to predict putative prophage regions. Phage_Finder (Fouts et al., 2006) is a standalone software program based on a heuristic algorithm to identify prophage regions in completely sequenced bacterial genomes. VirSorter (Roux et al., 2015) supports the detection of viral segments in microbiome sequencing data. PHASTER is a popular webserver for the identification and annotation of prophage sequences in prokaryotic genomes and plasmids (Wishart et al., 2016). Prophage Hunter (Xiao et al., 2019) provides a one-stop web service to identify prophage regions in bacterial genomes and evaluate the activity of the prophages. All these tools have substantially revolutionized the prediction of prophages in bacterial genomes. However, despite supporting prophage detection from massive bacterial genomes, Phage_Finder and VirSorter have limitations in speed and predictive power. PHASTER and Prophage Hunter support predictions using the webserver but cannot perform large-scale predictions of high-throughput microbiome sequencing data. Here, we introduce DBSCAN-SWA, a tool inspired by previous algorithms and tools, to detect prophages. DBSCAN-SWA has the fastest running speed and outperforms previous tools in detection rate and applicability for the prediction of complete and incomplete sequencing data (**Table 1**).

**Table 1.**
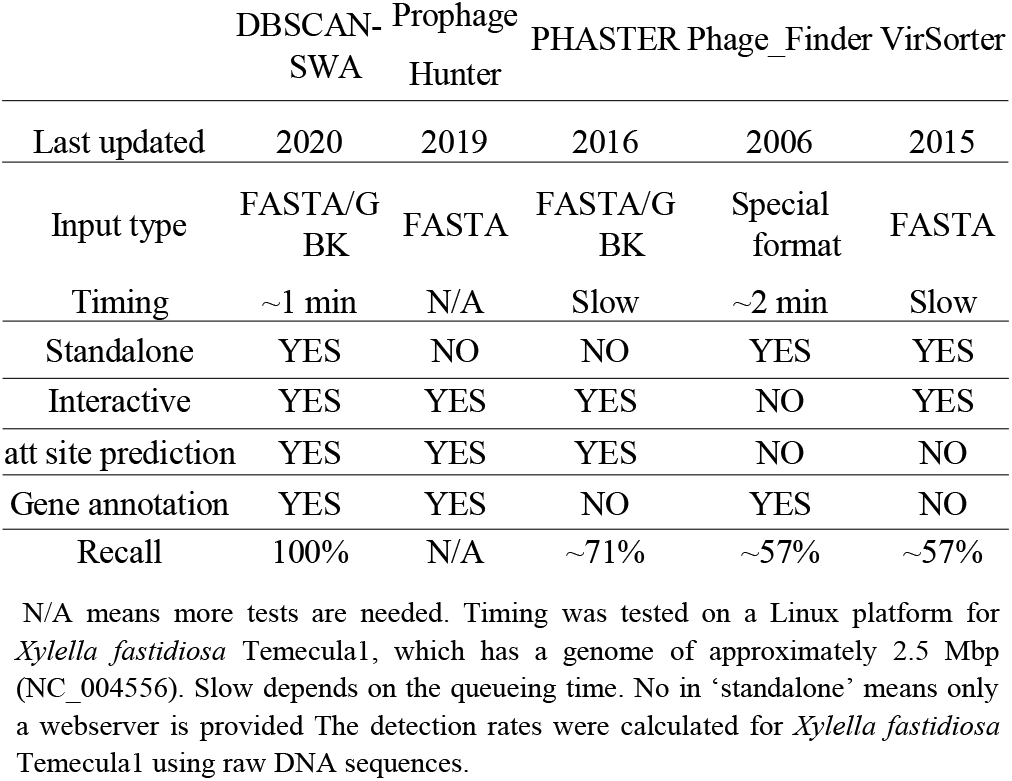
Comparison of DBSCAN-SWA with other prophage detection tools

## 2 Methods and Implementation

### 2.1 Prophage detection

DBSCAN-SWA implements an algorithm combining density-based spatial clustering of applications with noise (DBSCAN) and a sliding window algorithm (SWA) to detect putative prophages with reference to the theory underlying PHASTER, which is the most popular prophage detection tool but has no standalone version or source code available. Furthermore, we made some improvements. DBSCAN-SWA accepts two types of bacterial sequence files (multi-FASTA, GBK) for input. If a multi-FASTA input file is received, gene prediction and annotation will be performed by Prokka (Torsten, 2014) to obtain a standard GenBank format file with tRNA sites annotated using ARAGORN (Laslett et al., 2004). If a GenBank annotated file is submitted, gene annotations including protein sequences, descriptions and tRNA sites will be extracted for subsequent analysis. Considering prophage regions are composed of phage or phage-like genes clustered in a local region in the bacterial genome (Zhou et al., 2011), phage or phage-like proteins are first identified using Diamond BLASTP (e-values < 1e-7) (Buchfink et al., 2015) by searching against DBSCAN-SWA’s local viral UniProt TrEML reference database (UniProt C, 2014), which greatly improves the efficiency of DBSCAN-SWA compared to tools such as PHASTER using BLASTP. Then, DBSCAN is performed to detect phage protein clusters with the minimal number of phage-like genes required to form a prophage cluster (set to 6 proteins as the default parameter value) and the maximal spatial distance between two neighbor genes within the same cluster, which reflects the protein density within the prophage region (set to 3000 bps as the default parameter value). These two parameters are learned using a gradient method based on 184 manually curated prophage regions (Casjens et al., 2003) by trying the minimal prophage size from 6 to 10 proteins (step=1) and the protein density from 3000 to 10000 bp (step=1000 bp). Additionally, considering the biological features of prophages from different bacterial species may vary case by case, DBSCAN-SWA supports users to flexibly modify the two key parameters of DBSCAN, while PHASTER provides only fixed parameters. DBSCAN-SWA also uses SWA to scan specific key phage-related proteins in the GenBank file, such as ‘protease’, ‘integrase’, ‘transposase’, ‘terminase’, ‘lysis’, ‘bacteriocin’ and other key phage structural genes. Regions with at least six key proteins within a moving window of 60 proteins are considered putative prophage regions. The borders of the prophage region are determined as the positions of the first and the last occurred key proteins. Because the integrase enzyme typically encoded within temperate phages usually determines the specificity of integration sites (Williams KP, 2002), putative attachment sites will be examined for putative prophage clusters containing integrase. Using the integrase protein as an anchor, as each cluster contains an integrase, the sequences of 10 upstream and downstream proteins in the cluster will be extracted to detect the putative attL-attR pairs (mismatch=0) using BLASTN with the parameters ‘-task blastn-short −evalue 1000’. The attL-attR pair with the highest bit score and length>=12 bp is considered the putative att sites of the prophage region. Finally, each prophage region is assigned a taxonomy by a majority vote based on the detected phage-like genes with annotated taxonomic information within the prophage.

### 2.2 Phage annotation for prophage

DBSCAN-SWA provides two ways to annotate infecting phages for the predicted prophage regions. If a candidate phage genome is given, DBSCAN-SWA will evaluate the similarity between the integrated prophage(s) and the phage genome(s) based on three prophage-related features, defined to evaluate the similarity between the integrated prophage(s) and the phage genomes based on homologous protein alignment by Diamond BLASTP and nucleotide alignment by BLASTN (Supplementary Table S2). Alternatively, users can choose the local phage genome and protein database (PGPD) containing 10,463 complete phage genome sequences and 684,292 nonredundant phage proteins collected from millardlab (http://millardlab.org/bioinformatics/bacteriophage-genomes/) to predict the infecting phages by a Diamond BLASTP and a BLASTN search against PGPD.

## 3 Results

DBSCAN-SWA is an integrated tool for the detection of prophages that combines ORF prediction and gene function annotation, phage-like gene cluster detection, attachment site identification, and infecting phage annotation (Fig. 1A) with user-friendly outputs (Fig. 1B, 1C). To evaluate the performance of DBSCAN-SWA, 184 manually curated prophages from 50 complete bacterial genomes (Casjens et al., 2003) are collected to examine the prophage prediction capability based on recall (the number of correctly predicted prophages detected divided by the number of total prophages) and precision (the number of correctly predicted prophages divided by the number of total predicted prophages). The results show that DBSCAN-SWA performs the best with recalls of 84% or 85% for GenBank annotated sequences or raw DNA sequences, compared to PHASTER with recall of 86% or 82% and Phage_Finder and VirSorter with recall of 63% and 74% (Supplementary Table S1). With the best recall, DBSCAN-SWA shows a highly nonconservative predictive power with a precision of ~0.45. This means that DBSCAN-SWA fully considers the enrichment of viral genes in a putative prophage and will probably identify additional putative active prophages that can be induced by physiological cues. Moreover, DBSCAN-SWA presents better predictive power in NGS data than Phage_Finder and VirSorter based on the analysis of 19,989 contigs of 400 bacterial genomes in the human gastrointestinal tract collected from HMP (https://www.hmpdacc.org/hmp). DBSCAN-SWA predicts 2253 prophages on 1469 contigs from 389 bacterial genomes in approximately 13 hours with a detection rate (the percentage of bacterial genomes with putative prophages detected) of (389/400) 97%, while Phage_Finder predicts 580 prophages from 261 bacterial genomes in approximately 14 hours with a detection rate of (320/400) 80%. Compared to VirSorter, DBSCAN-SWA runs 6 times faster than VirSorter, which takes approximately 63 hours to predict 3016 prophages from 384 bacterial genomes (Supplementary Table S3, S4, S5). Simultaneously, DBSCAN-SWA also has a good degree of agreement with the prediction results of Phage_Finder, sharing 433 prophages with Phage_Finder (433/580=74.7%) and 1186 prophages with VirSorter (1186/3016=39.3%). It proves that DBSCN-SWA can predict putative prophages for increasingly high-throughput sequencing data and outperforms the existing standalone prophage prediction tools in terms of either efficiency or predictability. In summary, the highlights of DBSCAN-SWA are as follows:

- **High efficiency**. DBSCAN-SWA takes approximately 1.35 min~6.8 min to detect prophages in complete bacterial genomes (1.2 Mbp~7 Mbp).
- **High recall**. DBSCAN-SWA achieves an excellent recall of 100% for *Xylella fastidiosa* Temecula1 using raw DNA sequences, while PHASTER obtains only 71.4% (Fig. 1B).
- **Suitable for high-throughput sequencing data**. DBSCAN-SWA was compared to VirSorter and Phage_Finder, the only tools with standalone versions. DBSCAN-SWA showed better performance in efficiency, installation and usage than VirSorter, which is difficult to install based on a complex configuration environment, whereas DBSCAN-SWA is well packaged and easy to install. In addition, DBSCAN-SWA is suitable for both complete and incomplete sequenced genomes, while Phage_Finder is fit only for complete ones.
- **Provide phage annotation**. DBSCAN-SWA provides a custom phage database to facilitate the annotation of prophage regions. In addition, the similarity between the integrated prophage(s) and the phage genome(s) can be evaluated based on three prophage-related features (Fig. 1C).
- **User-friendly visualizations**. DBSCAN-SWA provides a userfriendly interactive HTML page to browse prophages in a genome viewer and detailed prophage information and bacterium-phage interactions in tables (Fig. 1 B, 1C).
- **Freely modified parameters**. DBSCAN-SWA enables users to adjust the parameters for phage-like protein identification, att site identification and phage annotation to meet their requirements for the prediction results based on their knowledge of prophages and phage-host interactions.

---

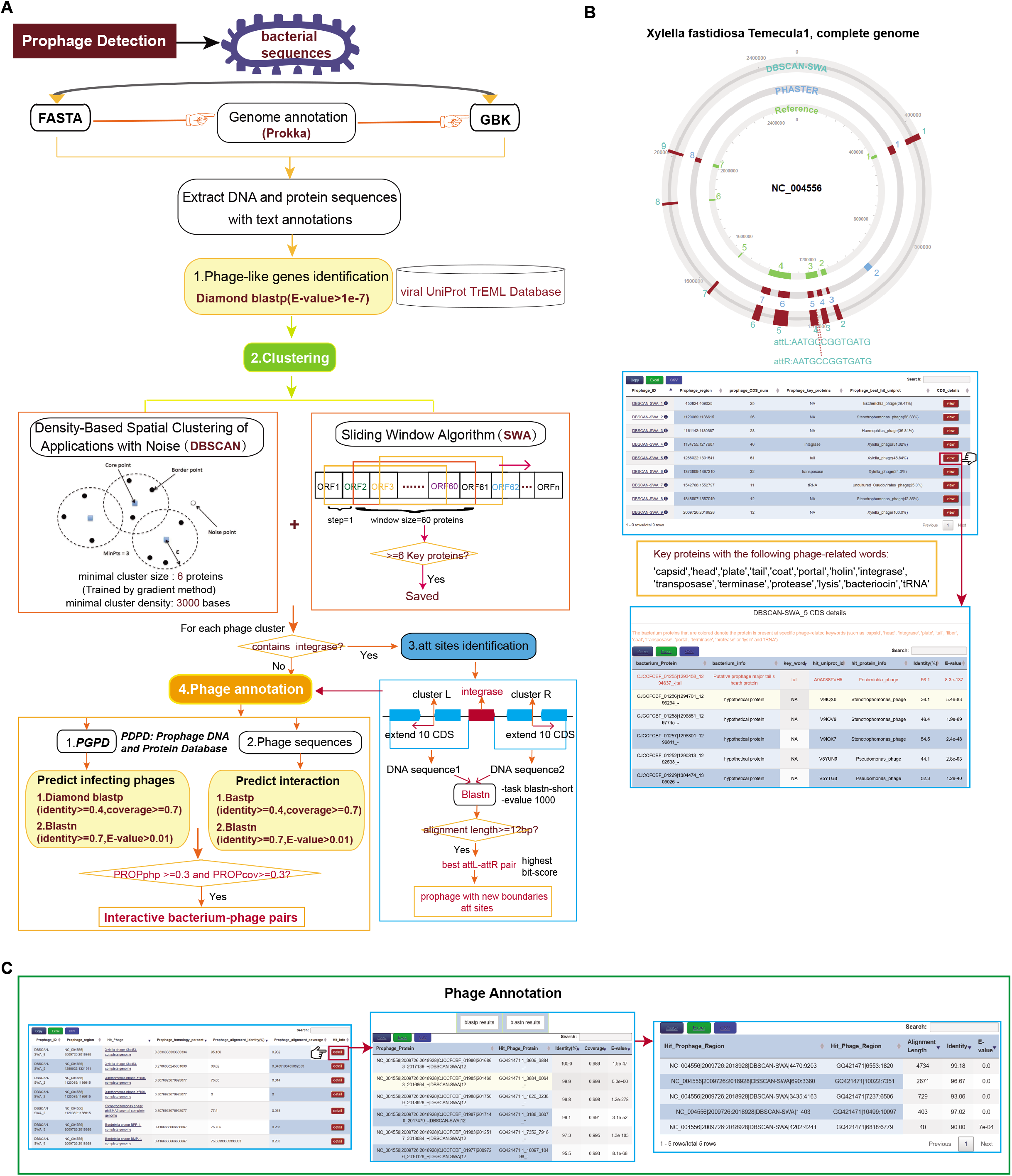

